# Replication-coupled chromatin assembly cooperates with ATR/CHK1 signaling to suppress DNA damage and premature MMB-dependent mitotic transcription

**DOI:** 10.64898/2026.09.27.754407

**Authors:** Devi Bala Murugan, Dörthe Gertzmann, Merima Ramic, Nils Greim, Aikaterini Boura, Karen M. Auweiler, Lea Damaschke, Christina Schülein-Völk, Carsten P. Ade, Martin Eilers, Stefan Gaubatz

**Affiliations:** Department of Biochemistry and Cell Biology, Theodor Boveri Institute, Biocenter, Julius Maximilian University Würzburg, Am Hubland, 97074 Würzburg, Germany; Core Unit High-Content Microscopy, Theodor Boveri Institute, Biocenter, Julius Maximilian University Würzburg, Am Hubland, 97074 Würzburg, Germany; Department of Biochemistry and Molecular Biology, Theodor Boveri Institute, Biocenter, Julius Maximilian University Würzburg, Am Hubland, 97074 Würzburg, Germany

## Abstract

Faithful genome duplication requires coordination of DNA replication, chromatin assembly, and checkpoint signaling to prevent inappropriate mitotic progression. Using a targeted siRNA screen for factors modulating CHK1 inhibitor-induced DNA damage and premature mitotic entry, we identified replication-coupled chromatin assembly as a determinant of checkpoint dependence. Acute depletion of the CAF-1 subunit CHAF1A modestly delayed S-phase progression but caused little DNA damage, whereas inhibition of CHK1 or ATR triggered extensive DNA damage, premature mitotic entry, and loss of viability. Transcriptomic analyses revealed that CHK1 inhibition induced a transcriptional response enriched for B-MYB/Myb-MuvB (MMB) target genes in CHAF1A-deficient cells. Notably, induction of these genes occurred without corresponding increases in promoter accessibility, suggesting that enhanced MMB-dependent transcription is largely independent of widespread promoter opening. B-MYB depletion attenuated MMB target-gene induction, premature mitotic entry, and DNA damage. Depletion of ASF1 and NAP1L1 similarly enhanced checkpoint inhibitor sensitivity, indicating that increased dependence on ATR-CHK1 signaling is a broader consequence of impaired histone chaperone function. Together, our findings establish a functional link between replication-coupled chromatin assembly, ATR-CHK1 signaling, and MMB-dependent mitotic transcription.

## INTRODUCTION

Faithful genome duplication during S phase requires coordinated DNA replication, chromatin assembly, and cell-cycle progression. Disruption of these processes can lead to replication stress, a hallmark of many cancers and typically associated with slowing or stalling of replication forks and increased genome instability ^1–3^. To maintain genome integrity, cells activate the ATR-CHK1 checkpoint pathway, which stabilizes replication forks, restrains origin firing, and prevents premature mitotic entry until DNA replication has been completed ^4–7^. Tumor cells can experience elevated levels of replication stress, making the ATR-CHK1 pathway a possible therapeutic target ^8,9^. Pharmacological inhibition of ATR or CHK1 induces DNA damage, unscheduled mitotic entry and chromosome segregation defects and shows selective activity in tumors with high replication stress ^10–12^. However, the molecular determinants that govern cellular sensitivity to checkpoint inhibition remain incompletely understood.

In addition to replication factors and checkpoint proteins, chromatin assembly pathways play essential roles during DNA replication. Newly synthesized DNA must be packaged into nucleosomes to preserve chromatin structure and genome stability ^13^. The chromatin assembly factor 1 (CAF-1) complex, composed of the subunits CHAF1A, CHAF1B, and RBBP4, is a replication-coupled histone H3-H4 chaperone that deposits newly synthesized histones onto nascent DNA ^14–19^. CAF-1 has also been implicated in heterochromatin maintenance, DNA repair, and cell-fate regulation, and loss of CAF-1 function has been associated with replication defects and genome instability ^20–26^. Recent work has demonstrated that acute CAF-1 depletion slows replication fork progression, prolongs S phase, and perturbs chromatin maturation ^27^.

Although CAF-1 is essential for replication-coupled chromatin assembly and normal replication dynamics, it remains unclear how defects in chromatin assembly influence cellular dependence on ATR-CHK1 checkpoint signaling. Moreover, transcriptional programs controlling cell-cycle progression can contribute to the cellular consequences of checkpoint abrogation. For example, the B-MYB/Myb-MuvB (MMB) multiprotein complex promotes expression of genes required for late S phase, G2 phase, and mitosis and has been implicated in cellular responses to replication stress after ATR-CHK1 inhibition ^28–30^.

Because most previous genetic screens have focused on long-term survival after checkpoint inhibition, less is known about chromatin regulators that influence the immediate cellular consequences of checkpoint failure ^28,31–36^. We therefore performed a targeted siRNA screen using acute CHK1 inhibitor-induced DNA damage and premature mitotic entry as functional readouts to identify chromatin pathways that modulate the early response to checkpoint abrogation. Using this approach, we identified replication-coupled histone chaperones as a determinant of cellular dependence on ATR-CHK1 signaling. Using inducible depletion systems, high-content microscopy, and transcriptomic analyses, we show that loss of CHAF1A enhances DNA damage, premature mitotic entry, and cell death following inhibition of ATR-CHK1 signaling. Mechanistically, CHK1 inhibition in CHAF1A-deficient cells induces a mitotic transcriptional response enriched for B-MYB/MMB target genes, and depletion of B-MYB attenuates these phenotypes. Together, our findings reveal a functional link between replication-coupled chromatin assembly, checkpoint signaling, and mitotic transcription.

## RESULTS

### An siRNA screen identifies CAF-1 as a protective factor during CHK1 inhibition

To identify chromatin regulators that influence the early cellular response to CHK1 inhibition, we previously performed a targeted siRNA screen against epigenetic and chromatin-associated genes in A549 cells ^37^. The screen quantified CHK1i-induced DNA damage and premature mitotic entry in S-phase cells and identified factors whose depletion either suppressed or enhanced these phenotypes (Figure 1A,B). In our previous study, we investigated suppressing hits and identified TRRAP as a coactivator of MMB-dependent mitotic transcription ^37^. Here we focused on sensitizing hits. Among the strongest sensitizers were several histone chaperones involved in replication-coupled chromatin assembly, including the core CAF-1 subunits CHAF1A and CHAF1B, as well as NAP1L1 (Figure 1B). Because CAF-1 is the major histone H3-H4 chaperone mediating replication-coupled nucleosome assembly, we focused on CHAF1A for further mechanistic investigation ^38^. Individual siRNAs targeting CHAF1A, CHAF1B or NAP1L1 each enhanced CHK1i-induced γH2AX accumulation and premature mitotic entry, validating the screening results (Figures 1C and S1A). Similarly, depletion of ASF1A, a histone H3-H4 chaperone upstream of CAF-1 and not represented in the original library ^38^, increased CHK1 inhibitor sensitivity (Figure S1B,C). Together, these findings indicate that impaired replication-coupled histone chaperone function increases cellular dependence on ATR-CHK1 signaling.

**Figure 1:**
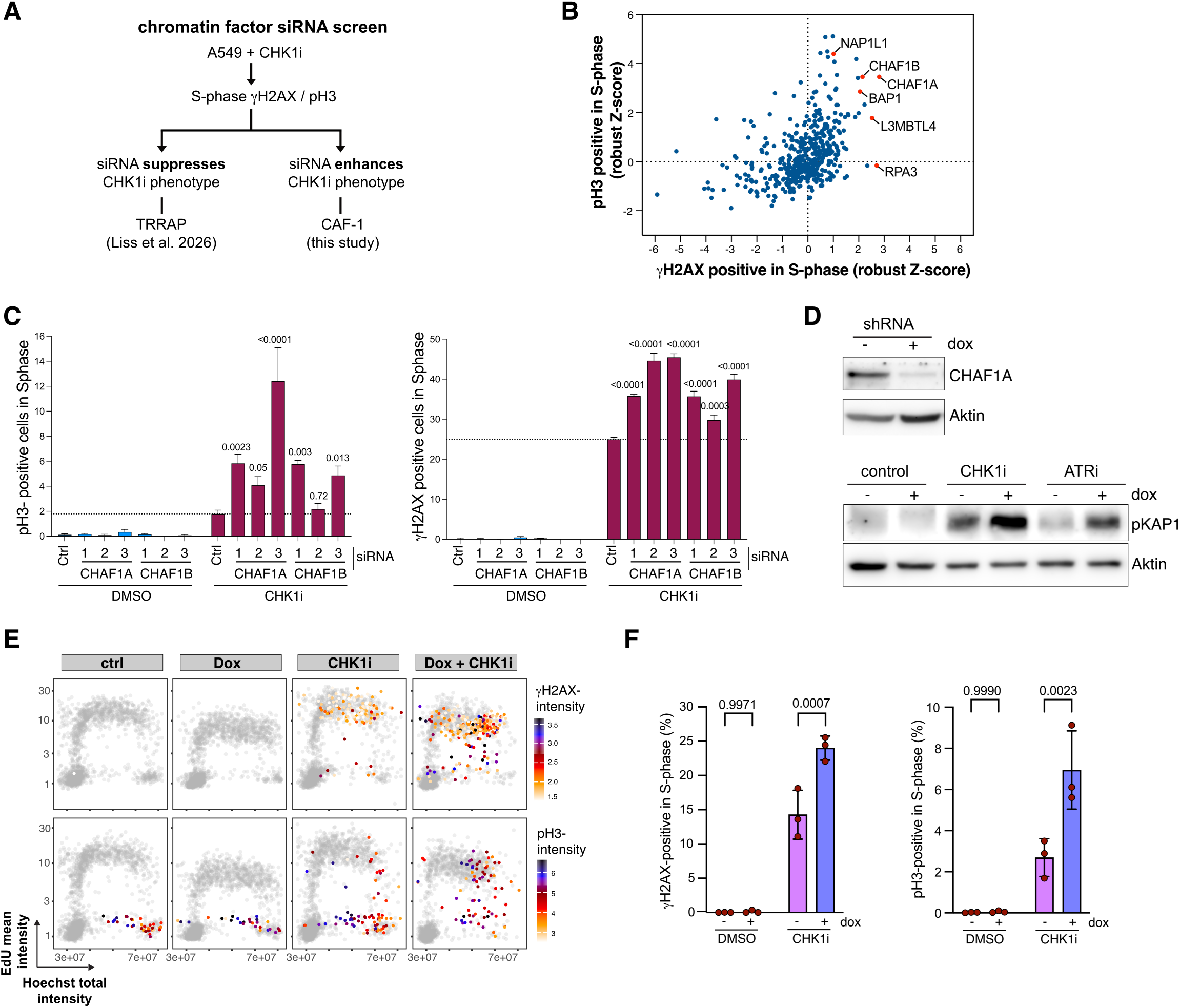
An siRNA screen identifies CAF-1 as a protective factor during CHK1 inhibition. A) Schematic of the targeted siRNA screen. CHK1i-induced DNA damage (γH2AX) and premature mitotic entry (pH3) were quantified in EdU-positive S-phase cells. Whereas siRNAs that suppress the phenotype were characterized previously ^37^, the present study focuses on sensitizing hits. B) Scatter plot of robust Z-scores for CHK1i-induced pH3 and γH2AX positivity in S-phase cells from the previously reported siRNA screen ^37^, highlighting the CAF-1 subunits CHAF1A and CHAF1B and the histone chaperone NAP1L1. C) Validation with three independent siRNAs targeting CHAF1A and CHAF1B. Percentage of cells positive for γH2AX and pH3 in S phase was quantified after treatment with CHK1i or DMSO. Data are shown as mean ± SEM. Two-way ANOVA (n = 3). D) Immunoblotting of A549 cells expressing a doxycycline-inducible CHAF1A-specific shRNA. pKAP1(S824) was used as a DNA damage signaling marker. Cells were treated with doxycycline for 48 h and with 100 nM prexasertib (CHK1i) or 5 µM ceralasertib (ATRi) for the final 4 h. E) High-content microscopy of cells treated with doxycycline and CHK1i as indicated. S-phase cells were identified by EdU incorporation. DNA content (x axis) and EdU incorporation (y axis) are shown for individual cells, color-coded according to mean γH2AX or pH3 fluorescence intensity. Representative experiment shown (n = 2,000 cells). F) Quantification of γH2AX- and pH3-positive S-phase cells from experiments shown in E). Data are shown as mean ± SD. Two-way ANOVA (n = 3).

To enable inducible knockdown of CHAF1A, we next generated A549 cells with a doxycycline-inducible shRNA targeting CHAF1A (Figure 1D). KAP1 phosphorylation at serine 824, a marker of ATM-dependent double-strand signaling, remained low upon CHAF1A depletion alone or after short-term CHK1 or ATR inhibition (Figure 1D). In contrast, robust KAP1 phosphorylation was detected when CHAF1A-depleted cells were treated with CHK1 or ATR inhibitors, confirming that CAF-1 loss increases DNA damage when checkpoint signaling is compromised. High-content microscopy further confirmed S-phase-specific DNA damage and premature mitosis upon simultaneous CHAF1A depletion and CHK1 inhibition (Figure 1E,F).

#### Acute CHAF1A degradation establishes an immediate requirement for checkpoint signaling during S phase

To distinguish the immediate consequences of CHAF1A loss from secondary effects of prolonged depletion, we generated CHAF1A mini auxin-inducible degron (mAID) cell lines. The mini auxin-inducible degron (mAID) cassette was knocked into the endogenous CHAF1A locus in A549 and HCT116 cells and the TIR1 F-box protein required for auxin-dependent degradation was stably expressed in these cells (Figure 2A) ^39^. Because prolonged CHAF1A depletion has previously been reported to induce a p53-dependent cell-cycle arrest ^39^, we additionally generated a CHAF1A-mAID HCT116 p53-knockout cell line. Treatment with the auxin analog 5-Ph-IAA led to near-complete CHAF1A depletion within six hours in these cell lines (Figure 2B).

**Figure 2:**
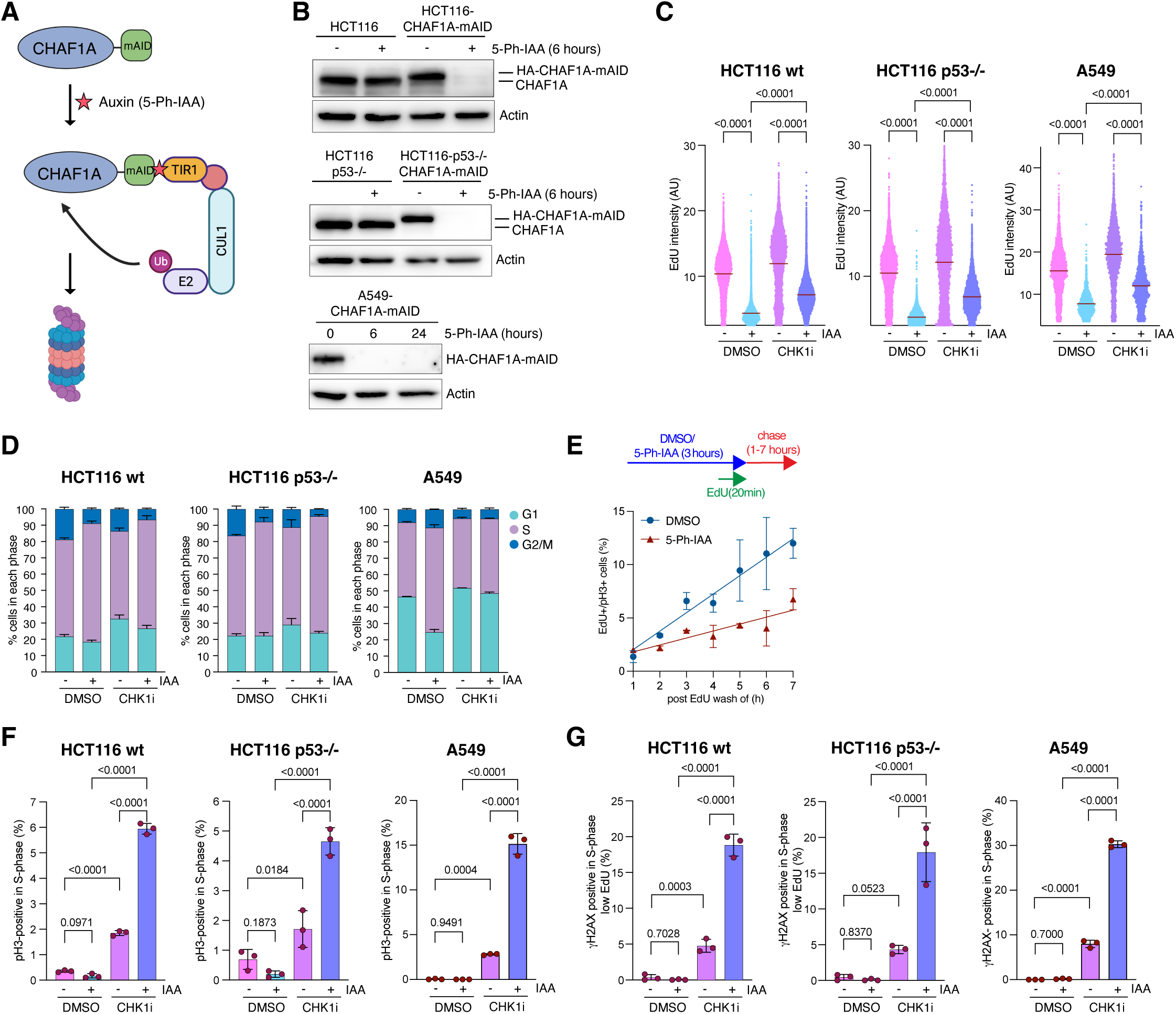
Acute CHAF1A degradation reveals an immediate requirement for checkpoint signaling during S phase. A) Schematic overview of auxin-inducible CHAF1A degradation. Cells were engineered to carry a mini auxin-inducible degron (mAID) fused to the endogenous CHAF1A locus and to express TIR1. Addition of the auxin analog 5-Ph-IAA induces rapid CHAF1A degradation. B) Immunoblot analysis confirming rapid CHAF1A degradation following 5-Ph-IAA treatment in HCT116 and A549 cells. Actin served as a loading control. C) Single-cell EdU incorporation analysis following CHAF1A degradation and CHK1 inhibition. Violin plots show EdU fluorescence intensities from a representative experiment (≥1697 cells analyzed per condition). HCT116 cells were treated with 5-Ph-IAA and CHK1i for 6 h as indicated. A549 cells were first treated with 5-Ph-IAA for 22 h followed by CHK1i treatment for 2 h. D) Cell-cycle distribution determined by high-content microscopy after treatment as in C). Data represent mean ± SD from three independent experiments. E) Pulse-chase analysis of S phase progression into mitosis. Cells were pulse-labeled with EdU for 20 min, chased for the indicated times, and stained for phospho-histone H3 (pH3). The fraction of EdU-positive cells entering mitosis (pH3-positive) is plotted as a function of duration of the chase. Data represent two independent experiments. F,G) Quantification of γH2AX-positive F) and pH3-positive G) S-phase cells following CHAF1A degradation and CHK1 inhibition. Treatments were performed as in C). Data represent mean ± SD from three independent experiments. Statistical significance was determined by one-way ANOVA.

Acute CHAF1A degradation reduced EdU incorporation, delayed S-phase progression and modestly increased the S-phase population, consistent with impaired replication dynamics (Figure 2C-E) ^26,27,40^. Consistent with increased origin firing following checkpoint abrogation, CHK1 inhibition partially restored EdU incorporation (Figure 2C) ^41–43^. Although CHAF1A depletion alone produced little γH2AX or pH3 accumulation, CHK1 inhibition strongly enhanced both phenotypes (Figure 2F,G). In HCT116 cells, γH2AX accumulated preferentially in S-phase cells with low EdU incorporation (Figure S2C). 5-Ph-IAA treatment of TIR1-expressing control HCT116 and A549 cells without the CHAF1A-mAID allele did not enhance γH2AX or pH3 staining after CHK1i treatment, indicating that the observed phenotypes were specifically caused by CHAF1A degradation (Figure S2A,B).

To assess whether CHAF1A loss induces replication stress, we quantified FANCD2 intensity in S-phase cells after acute CHAF1A degradation and CHK1 inhibition ^41,42^. CHAF1A depletion by itself caused little increase in FANCD2 staining, whereas combined CHAF1A depletion and CHK1 inhibition markedly increased FANCD2 intensity in S-phase cells (Figure S2D). These findings indicate that defective chromatin assembly results in replication stress primarily when checkpoint signaling is compromised.

### CAF-1 deficiency creates a selective dependence on ATR-CHK1-WEE1 signaling

To determine whether the enhanced DNA damage and premature mitotic entry observed in CHAF1A-deficient cells affect long-term cell survival, we next performed clonogenic survival assays. ATR, CHK1, and WEE1 inhibitors had only modest effects on colony formation in control HCT116 cells but strongly impaired colony formation following CHAF1A depletion, indicating enhanced sensitivity to checkpoint inhibition following CAF-1 loss (Figure 3A,B). This sensitization was observed in both wild-type and p53-knockout HCT116 cells, showing that the interaction is largely independent of the p53 status. In contrast, CHAF1A depletion did not similarly sensitize cells to treatment with doxorubicin, a topoisomerase II inhibitor, camptothecin, a topoisomerase I inhibitor, or the CDK4/6 inhibitor palbociclib (Figure 3C,D). This suggests that CAF-1 loss does not generally increase sensitivity to all antiproliferative agents but preferentially enhances vulnerability to S-phase checkpoint inhibition.

**Figure 3:**
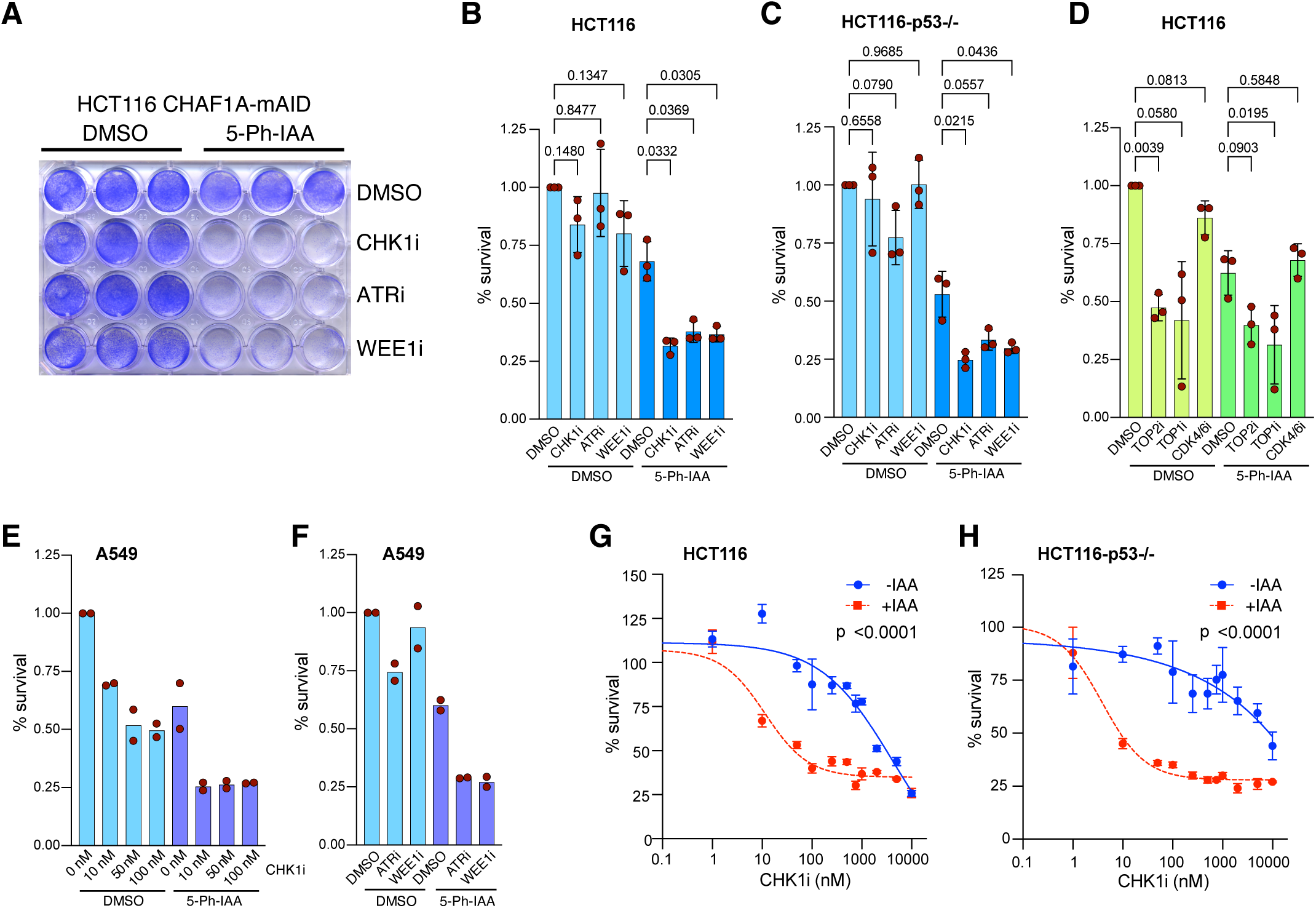
CAF-1 loss creates selective sensitivity to ATR, CHK1, and WEE1 inhibition. A) HCT116-CHAF1A-mAID cells were treated with DMSO or 5-Ph-IAA and with CHK1i, ATRi or WEE1i for 2 days. Colonies were fixed and stained with crystal violet. B)-F) Quantification of colony formation assays in the indicated cell lines after treatment with DMSO or 5-Ph-IAA and the indicated drugs for 48 hours (n = 3, one-way ANOVA). G) and H) Cell viability of HCT116-CHAF1A-mAID cells (wt and p53-/-) exposed to different concentrations of CHK1i for 48 h (n = 3, extra sum-of-squares F test).

A549 cells showed a similar pattern, with combined CHAF1A depletion and ATR, CHK1, or WEE1 inhibition causing a reduction in clonogenic survival compared with either treatment alone (Figure 3E,F). Consistent with these findings, viability assays using increasing concentrations of the CHK1 inhibitor prexasertib confirmed increased sensitivity of both wild-type and p53-knockout HCT116 cells to CHK1 inhibition following CHAF1A depletion (Figure 3G,H).

#### CHK1 inhibition induces MMB-associated mitotic transcription in CAF-1-deficient cells

To identify mechanisms underlying checkpoint dependence following CAF-1 loss, we performed RNA sequencing after CHAF1A depletion, short-term CHK1 inhibition, or the combination of both treatments. CHAF1A depletion for 24 h induced widespread transcriptional changes characterized predominantly by gene upregulation (Figure 4A). In contrast, CHK1 inhibition for 2 h had only modest effects on gene expression (Figure S3A), while the transcriptional changes after combined treatment were largely dominated by CHAF1A depletion (Figure S3B). We therefore directly compared CHAF1A-depleted cells treated with CHK1 inhibitor to CHAF1A-depleted cells alone to identify transcriptional responses specifically associated with checkpoint inhibition in cells without functional CAF-1 (Figure 4B). This analysis identified a discrete set of differentially expressed genes enriched for cell-cycle regulators, including genes involved in mitotic entry and progression such as KIF20A, PLK1, and FAM83D (Figure 4B,C). The limited number of differentially expressed genes is consistent with the short duration of CHK1 inhibitor treatment. Gene set enrichment analysis revealed significant enrichment of mitotic and Myb-MuvB (MMB)-associated transcriptional programs among genes induced by CHK1 inhibition in CHAF1A-depleted cells (Figure 4D, Figure S3C). A subset of MMB target genes was more strongly induced by CHK1 inhibition in CHAF1A-depleted cells than in control cells (Figure 4C,E). Increased expression of selected MMB target genes was confirmed by RT-qPCR (Figure 4F). RT-qPCR also validated induction of DHRS2, a transcriptional target of CHAF1A depletion (Figure 4F). Together, these findings indicate that CHK1 inhibition in CAF-1-deficient cells induces a mitotic transcriptional response enriched for MMB target genes.

**Figure 4:**
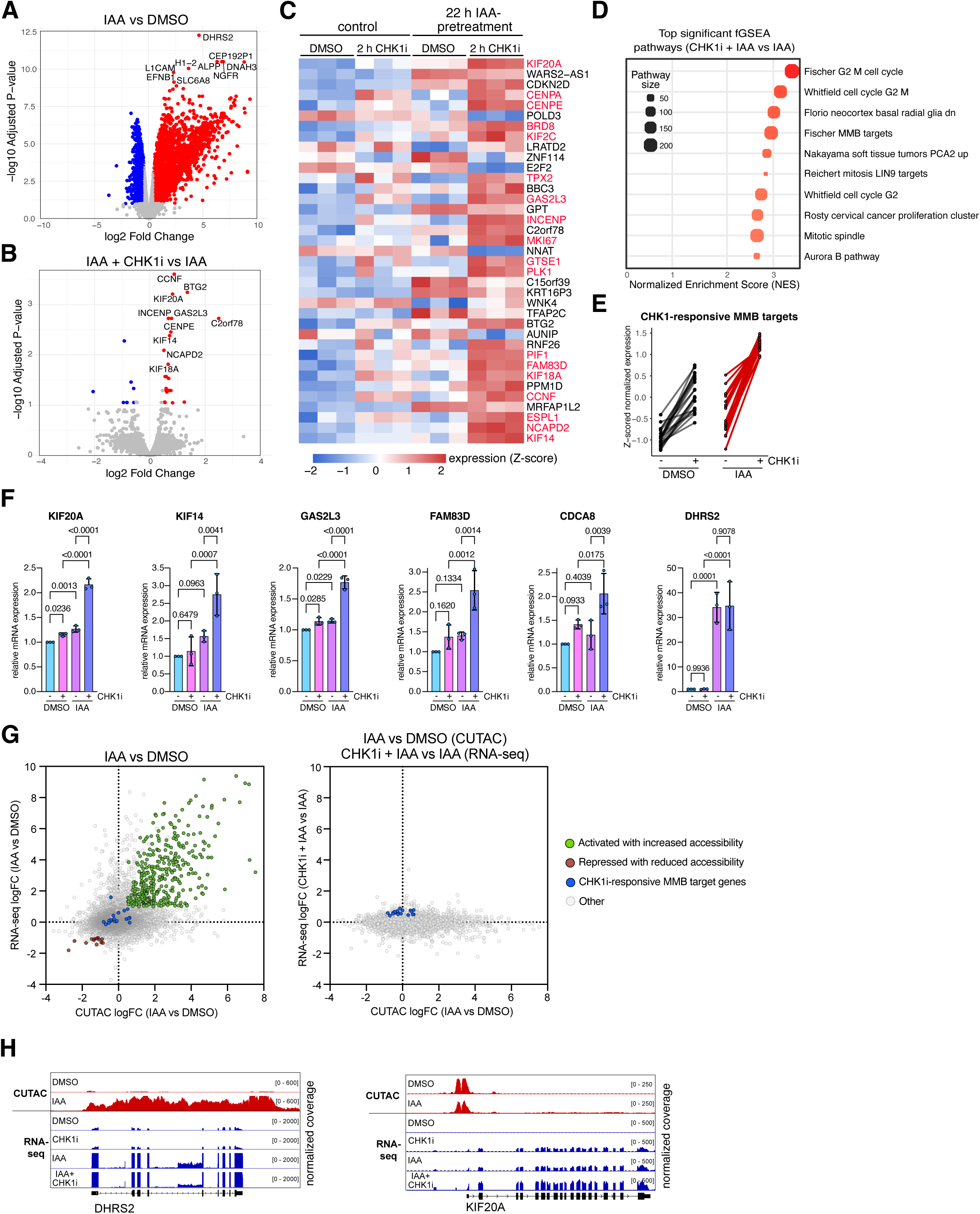
CHK1 inhibition induces MMB-associated mitotic transcription in CAF-1-deficient cells. A, B) Volcano plots of differentially expressed genes following 24 h 5-Ph-IAA treatment (A) or CHK1 inhibition in 5-Ph-IAA-treated cells (B). Significantly regulated genes (absolute log2 fold change > 0.5 and adjusted P < 0.1) are highlighted in red (upregulated) or blue (downregulated). C) Heatmap showing the top differentially expressed genes following CHK1 inhibition in CHAF1A-depleted cells. MMB target genes are highlighted in red. Data represent three biological replicates (n = 3). D) Gene set enrichment analysis (fGSEA) of genes differentially expressed following CHK1 inhibition in CHAF1A-depleted cells. The top enriched pathways ranked by normalized enrichment score (NES) are shown. E) Expression of MMB target genes significantly induced by CHK1 inhibition in CHAF1A-depleted cells (IAA+CHK1i versus IAA; FDR < 0.1; n = 20 genes). Normalized RNA-seq expression values are shown as row z-scores. Lines connect values for individual genes within the control (DMSO versus CHK1i) and CHAF1A-depleted (IAA versus IAA+CHK1i) conditions. F) RT-qPCR analysis of selected MMB target genes and the non MMB target gene DHRS2 following CHK1 inhibition, CHAF1A depletion, or combined treatment. Data are shown as mean ± SD (n = 3 independent replicates). Statistical significance was assessed by one-way ANOVA. G) Relationship between promoter accessibility and gene expression following CHAF1A depletion and checkpoint inhibition. Left: Scatter plot showing RNA-seq log2 fold changes and corresponding changes in promoter-associated CUTAC signal within ±1 kb of the transcription start site following CHAF1A depletion (IAA versus DMSO). Right: Cross-comparison of promoter accessibility changes following CHAF1A depletion (IAA versus DMSO) with RNA-seq expression changes induced by CHK1 inhibition in CHAF1A-depleted cells (IAA+CHK1i versus IAA). Each point represents one gene. Genes showing concordant increases in gene expression and promoter accessibility are highlighted in green, whereas genes showing concordant decreases in gene expression and promoter accessibility are highlighted in red. CHK1i-responsive MMB target genes defined in (E) are highlighted in blue. H) Genome browser tracks of the DHRS2 and KIF20A loci showing CUTAC signal and RNA-seq coverage under the indicated treatment conditions.

Given the established role of CAF-1 in replication-coupled chromatin assembly, we asked whether transcriptional responses following CHAF1A depletion and CHK1 inhibition could be explained by changes in promoter accessibility. We therefore performed CUTAC-based accessibility profiling and integrated promoter accessibility with RNA-seq across treatment conditions. Consistent with the established role of CAF-1 in chromatin assembly, transcriptional activation following CHAF1A depletion was frequently associated with increased promoter accessibility (Figure 4G, left panel). We next asked whether the chromatin accessibility state established by CHAF1A depletion predicted the subsequent transcriptional response to CHK1 inhibition. To address this, we compared promoter accessibility changes following CHAF1A depletion (IAA versus DMSO) with RNA expression changes induced by CHK1 inhibition specifically in CHAF1A-depleted cells (IAA+CHK1i versus IAA). Promoter accessibility changes induced by CHAF1A depletion showed little relationship with the subsequent transcriptional response to CHK1 inhibition (Figure 4G, right panel). Notably, MMB target genes that were induced following CHK1 inhibition did not show a corresponding tendency toward increased promoter accessibility following CHAF1A depletion. Thus, induction of the CHK1i-responsive MMB target genes is unlikely to result simply from prior local promoter opening caused by CAF-1 loss.

Consistent with this interpretation, acute CHK1 inhibition produced little additional promoter accessibility change in CHAF1A-depleted cells despite induction of a discrete transcriptional response (Figure S3D). Likewise, the combined IAA+CHK1i versus DMSO comparison largely recapitulated the accessibility-associated changes caused by CHAF1A depletion alone. Together, these findings suggest that CAF-1 loss enhances induction of a subset of MMB-regulated mitotic genes following checkpoint inhibition through a mechanism that cannot be readily explained by local promoter opening.

Representative genome browser tracks illustrate these different regulatory modes: DHRS2, a known transcriptional target of CAF-1 loss, showed coordinated increases in promoter accessibility and gene expression after CHAF1A depletion (Figure 4H). In contrast, the MMB target gene KIF20A was induced following CHK1 inhibition in CHAF1A-deficient cells without detectable changes in promoter accessibility. These findings suggest that induction of the CHK1i-responsive MMB target genes is more likely driven through altered transcription factor activity than through widespread changes in promoter accessibility.

### B-MYB contributes to checkpoint inhibitor sensitivity following CHAF1A loss

To determine whether B-MYB contributes to checkpoint dependence following CHAF1A loss, we generated CHAF1A-mAID A549 cells carrying a doxycycline-inducible shRNA specific for B-MYB and confirmed efficient depletion by immunoblotting (Figure 5A). B-MYB depletion significantly reduced DNA damage and premature mitotic entry following combined CHAF1A depletion and CHK1 inhibition (Figure 5B). Although depletion of B-MYB also attenuated responses to CHK1 inhibition alone, suppression of the combined phenotype indicates that B-MYB activity contributes to the increased checkpoint dependence of CHAF1A-deficient cells. Similar results were obtained following siRNA-mediated depletion of B-MYB in CHAF1A-mAID HCT116 cells, demonstrating that this requirement is not restricted to a single cell line (Figure S4A,B).

**Figure 5:**
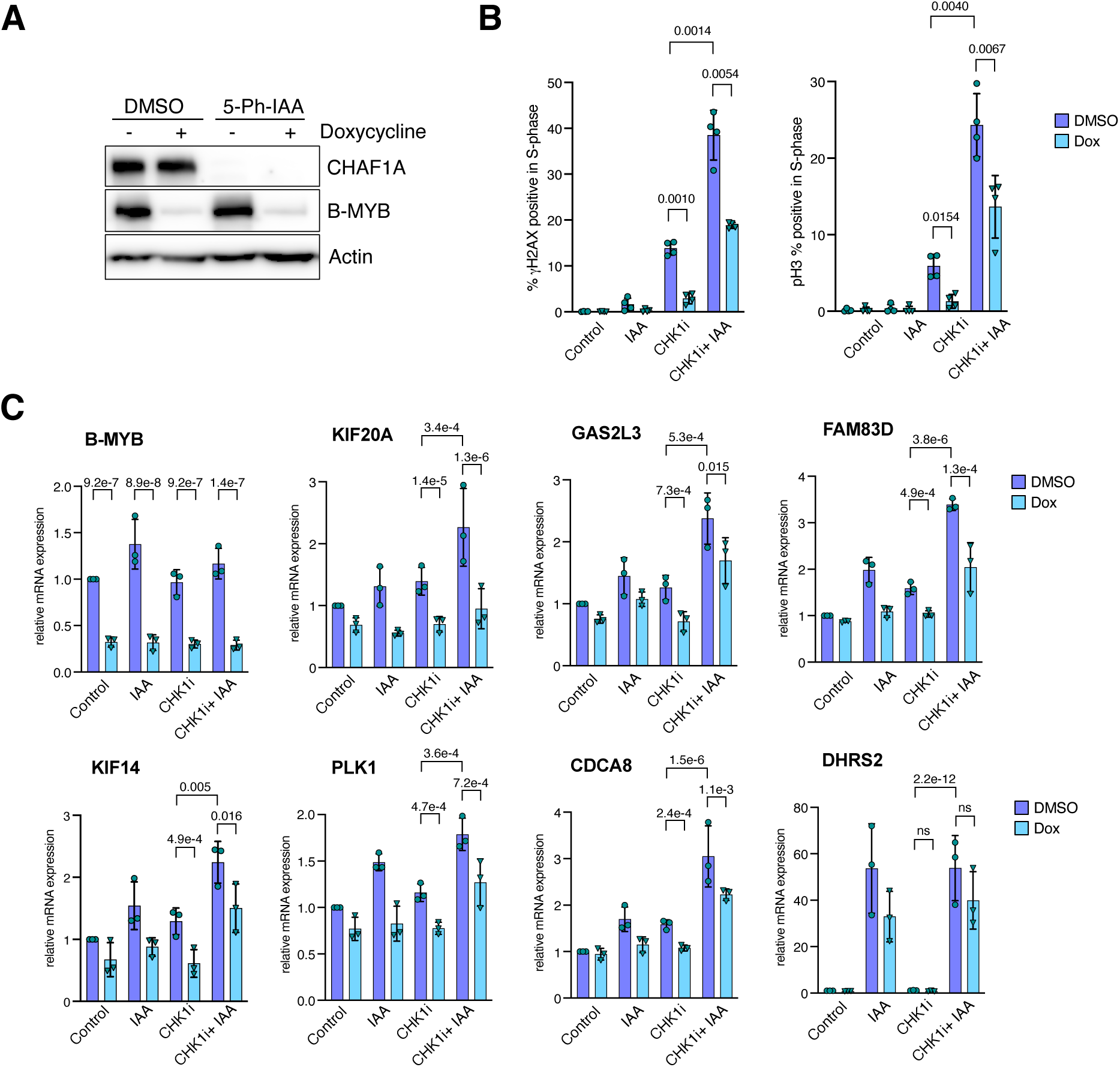
B-MYB contributes to CHK1i sensitivity following CHAF1A loss. A) Immunoblot analysis confirming doxycycline-induced depletion of B-MYB and 5-Ph-IAA-induced degradation of CHAF1A in A549 CHAF1A-mAID cells expressing a doxycycline-inducible shRNA targeting B-MYB. Actin served as a loading control. B) Quantification of the percentage of γH2AX-positive and phospho-histone H3 (pH3)-positive S-phase cells following CHAF1A depletion, CHK1 inhibition, and B-MYB depletion as indicated. Data represent mean and SD from four independent experiments. Statistical significance was determined by one-way ANOVA. C) RT-qPCR analysis of selected MMB target genes following CHAF1A depletion, CHK1 inhibition, and B-MYB depletion as indicated. Data represent mean ± SD from three independent experiments. Statistical significance was determined by one-way ANOVA.

To determine whether B-MYB is also required for the transcriptional response, we quantified expression of selected target genes by RT-qPCR. Efficient depletion of B-MYB mRNA was confirmed under all treatment conditions (Figure 5C). B-MYB depletion attenuated induction of representative MMB target genes following combined CHAF1A depletion and CHK1 inhibition, whereas expression of DHRS2, a CHAF1A-regulated gene that is not an MMB target, was unaffected (Figure 5C). These findings demonstrate that induction of these MMB target genes requires B-MYB. Notably, CHAF1A depletion did not alter B-MYB mRNA or protein abundance (Figure 5A,C), indicating that induction of the MMB transcriptional response is not driven by increased B-MYB expression.

Because B-MYB/MMB and CDK1 cooperate through positive feedback to drive mitotic entry ^28^, we next tested whether CDK1 activity contributes to checkpoint dependence following CHAF1A loss. Partial inhibition of CDK1 with RO-3306 reduced DNA damage and premature mitotic entry following combined CHAF1A depletion and CHK1 inhibition (Figure S4C). These findings support a model in which inappropriate activation of the B-MYB/MMB-CDK1 mitotic network contributes to the increased checkpoint dependence of CHAF1A-deficient cells.

## DISCUSSION

In this study, we identify replication-coupled chromatin assembly as a determinant of cellular dependence on ATR-CHK1 signaling. Loss of CAF-1 sensitized cells to ATR, CHK1, and WEE1 inhibition, and this phenotype was associated with activation of a subset of MMB target genes (summarized in Figure 6).

**Figure 6:**
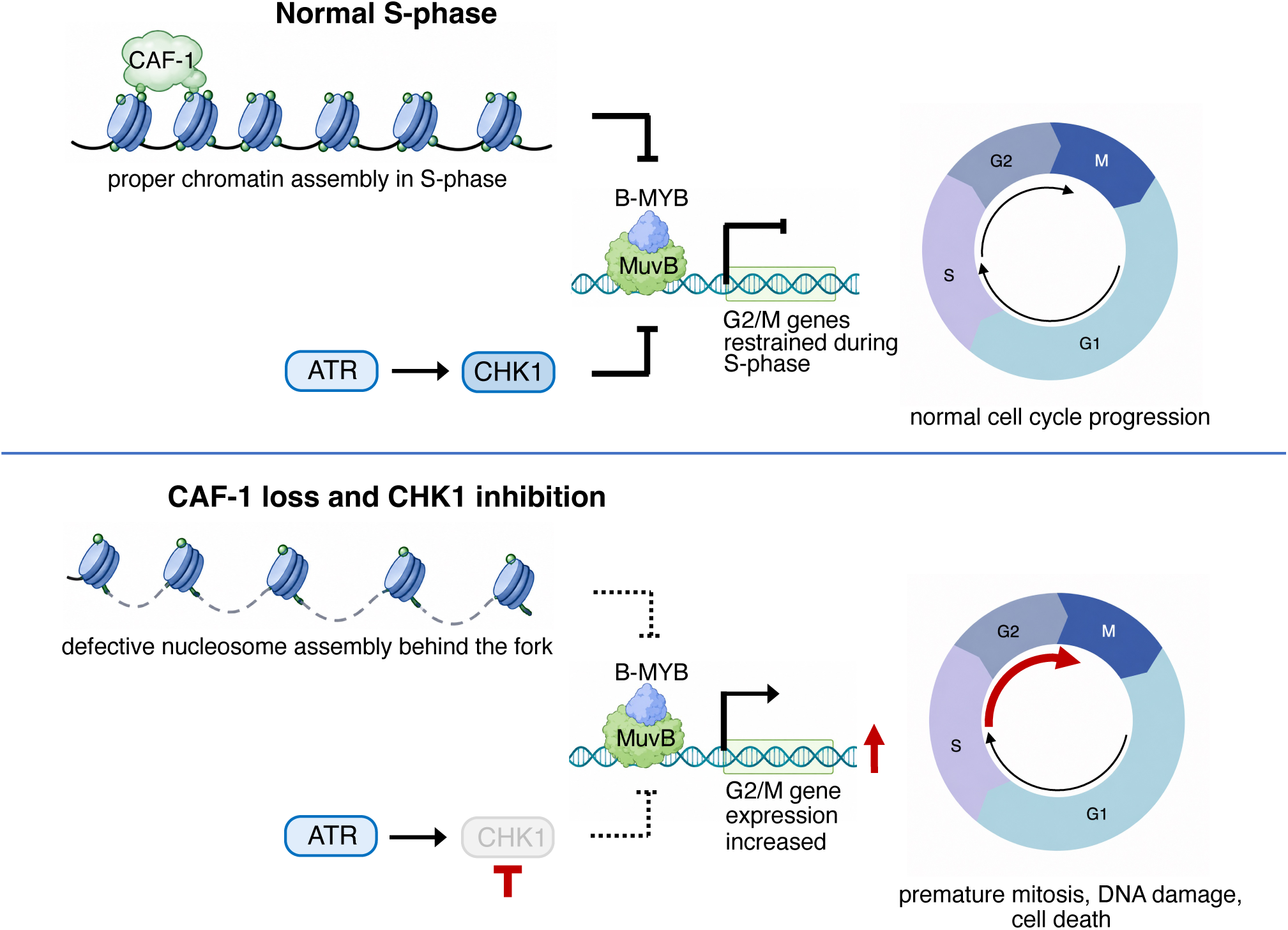
Model for enhanced checkpoint inhibitor sensitivity following CAF-1 loss. Under unperturbed conditions, replication-coupled chromatin assembly by CAF-1 and ATR– CHK1 signaling promote faithful S phase progression and prevent premature mitotic entry (top). Loss of CAF-1 alone causes a modest delay in S phase, but little DNA damage. However, when ATR or CHK1 signaling is inhibited in CAF-1-deficient cells, inappropriate B-MYB/MMB-dependent mitotic transcription contributes to premature mitotic entry, S phase-specific DNA damage, and loss of viability (bottom).

Importantly, CHAF1A depletion alone caused little DNA damage, indicating that acute CAF-1 loss does not simply trigger widespread genome instability. Earlier studies using dominant-negative CAF-1 demonstrated that disruption of S-phase chromatin assembly can induce DNA damage, checkpoint activation, and S-phase arrest ^43^. More recent work using acute CAF-1 depletion showed immediate slowing of replication fork progression and perturbation of nascent chromatin assembly without triggering a robust canonical ATR-CHK1 DNA damage response ^27^. Consistent with the latter findings, acute CHAF1A depletion in our system reduced EdU incorporation, modestly increased the fraction of cells in S phase, and delayed progression from S phase into mitosis.

A key finding of this work is that CHK1 inhibition in CHAF1A-deficient cells induces a mitotic transcriptional response enriched for B-MYB/MMB target genes. Previous work identified an intrinsic ATR-CHK1-dependent S/G2 checkpoint that restrains premature activation of the B-MYB/FOXM1 mitotic transcriptional program during DNA replication ^7,28^. Consistent with this model, CHK1 inhibition also induced MMB target genes in control cells, while a subset of MMB targets showed stronger induction after CHAF1A depletion. CUTAC analysis further showed that these CHK1i-responsive MMB target genes were not preferentially associated with increased promoter accessibility following CHAF1A depletion. This is notable because CHAF1A depletion caused substantial accessibility changes at other promoters that frequently correlated with transcriptional changes, suggesting that the enhanced MMB-dependent transcriptional response is largely independent of widespread promoter opening.

Functionally, B-MYB depletion attenuated MMB target-gene induction, CHK1 inhibitor-induced DNA damage, and premature mitotic entry, indicating that activation of the MMB transcriptional network contributes to checkpoint inhibitor sensitivity in CHAF1A-deficient cells. Because B-MYB depletion also attenuated the response to CHK1 inhibition in control cells, B-MYB is unlikely to function exclusively downstream of CAF-1 deficiency. Rather, our data support a model in which replication-coupled chromatin assembly cooperates with ATR-CHK1 signaling to restrain inappropriate activation of the B-MYB/MMB mitotic transcriptional program during S phase. Consistent with this model, partial CDK1 inhibition reduced both DNA damage and premature mitotic entry following CHK1 inhibition, supporting the idea that inappropriate activation of the B-MYB/MMB-CDK1 network contributes to checkpoint inhibitor sensitivity when chromatin assembly is impaired.

The identification of NAP1L1 and ASF1 as additional histone chaperones whose depletion enhances CHK1 inhibitor-induced phenotypes suggests that checkpoint dependence is a broader consequence of defective histone deposition and chromatin assembly. CAF-1, ASF1, and NAP1L1 act at distinct but interconnected steps of nucleosome assembly ^13^. Thus, perturbation of multiple histone chaperone pathways may increase dependence on ATR-CHK1 signaling to prevent DNA damage and inappropriate mitotic progression during S phase.

Our data do not establish how defective chromatin assembly influences the MMB-dependent transcriptional response to checkpoint inhibition. The absence of widespread promoter opening at CHK1i-responsive MMB target genes suggests that other consequences of defective chromatin assembly may influence MMB activity or subsequent transcriptional activation. Given that B-MYB/MMB activity is regulated through CDK-dependent phosphorylation and interactions with transcriptional cofactors, defining the upstream events linking defective chromatin assembly to mitotic transcription will require further study ^44^.

Together, our findings reveal a functional cooperation between replication-coupled chromatin assembly and ATR-CHK1 signaling that protects S-phase cells from DNA damage and inappropriate mitotic progression. They further establish MMB-dependent mitotic transcription as a functional contributor to checkpoint inhibitor sensitivity following CAF-1 loss and suggest that defects in histone chaperone pathways may create vulnerabilities to ATR or CHK1 inhibition.

## METHODS

### Cell lines

A549, HCT116 and HCT116 p53-/-cells were cultured in RPMI medium supplemented with 10% FCS and 1% penicillin/streptomycin. Cells were treated with prexasertib (CHK1 inhibitor), ceralasertib (AZD6738, ATR inhibitor), adavosertib (AZD1775, WEE1 inhibitor), 5-Ph-IAA, camptothecin (topoisomerase I inhibitor), doxorubicin (topoisomerase II inhibitor), palbociclib (CDK4/6 inhibitor) and RO-3306 (CDK1 inhibitor) as indicated.

### Cloning

A lentiviral shRNA construct was generated by insertion of an shRNA specific for CHAF1A into pINDUCER10. For degron tagging of endogenous CHAF1A, homology arms flanking the CHAF1A C-terminus were cloned into a pJET vector containing a knock-in cassette with a blasticidin resistance gene, an HA epitope and a mini auxin-inducible degron tag (mAID) ^45^. To construct a CRISPR/Cas9 plasmid expressing a sgRNA targeting the C-terminus of CHAF1A, annealed oligonucleotides with the target sequence were cloned into pX458 (Addgene #48138).

### Stable cell line generation

To generate A549 and HCT116 cell lines with mAID-tagged CHAF1A, cells were co-transfected with the donor construct and pX458-gRNA-CHAF1A expressing a guide RNA targeting the CHAF1A C-terminus. Cells were selected with 10 µg/ml blasticidin S (Invivogen) and single clones were isolated. Clones with a homozygous knock-in were identified by genomic PCR. Homozygous clones were subsequently transduced with lentiviral TIR1-F74G (pRRL-SFFV-TIR1-F74G ^46^) and selected with hygromycin B (Sigma Aldrich).

### Lentiviral production and infection

Lentiviral particles were produced in HEK-293-T cells by co-transfecting the lentiviral vector with psPAX2 and pCMV-VSV-G. Virus supernatant was collected 48 h after transfection, filtered and diluted 1:1 with culture medium with 4 µg/ml polybrene (Sigma). Infected cells were selected 48 h after infection with the appropriate antibiotics for 7 days.

### Immunoblotting

Cells were lysed in TNN (50 mM Tris (pH 7.5), 120 mM NaCl, 5 mM EDTA, 0.5% NP40, 10 mM Na4P_2_O_7_, 2 mM Na_3_VO_4_, 100 mM NaF, 1 mM PMSF, 1 mM DTT, 10 mM β-glycerophosphate, protease inhibitor cocktail (Sigma)). Proteins were separated by SDS-PAGE, transferred to a PVDF membrane and detected by immunoblotting with the primary and secondary antibodies. Antibodies are listed in Supplementary Table S1.

### siRNA transfection and siRNA screen

For individual knockdown experiments, siRNAs were purchased from Thermo Fisher Scientific and transfected using Lipofectamine RNAiMAX at a final concentration of 5 nM. siRNA sequences are listed in Table S1. The targeted siRNA screen and its analysis were performed previously ^37^, using the Silencer Select Human Epigenetics siRNA Library (Thermo Fisher Scientific, A30085). Briefly, A549 cells were reverse-transfected with pools of three siRNAs targeting 521 epigenetic and chromatin-associated genes and, 46 h later, treated with 100 nM prexasertib for 2 h. Cells were pulse-labeled with EdU and analyzed by high-content microscopy for γH2AX and pH3 within the EdU-positive population. The robust Z-scores are derived from the previously reported screen ^37^.

### RT-qPCR

Total RNA was isolated using Trizol (Thermo Fisher Scientific). cDNA was synthesized using random primers and RevertAid reverse transcriptase (Thermo Fisher Scientific). Quantitative real-time PCR was performed with the ABsolute QPCR SYBR Green Mix from Thermo Fisher Scientific using a qTower3G (Analytik Jena). Expression differences were calculated as described before ^47^. Primer sequences are listed in Table S1.

### RNA sequencing

Total RNA was isolated from three biological replicates. DNA libraries were generated using 1 µg RNA with the magnetic mRNA isolation module and NEBNext Ultra II RNA Library Prep Kit for Illumina (New England Biolabs). DNA libraries were amplified by PCR and quality was analyzed using the fragment analyzer (Advanced Analytical). Libraries were sequenced on the NextSeq 2000 platform (Illumina) (paired-end, 2 x 60 bp).

### CUTAC

Chromatin accessibility profiling was performed using CUTAC as described previously ^48^. Briefly, cells were harvested by trypsinization and 100,000 cells were bound to concanavalin A-coated magnetic beads for 10 min at room temperature. Following washes with Triton-Wash buffer (20 mM HEPES pH 7.5, 150 mM NaCl, 0.5 mM spermidine, 0.1% Triton X-100, protease inhibitor cocktail (Sigma, P8340, 1:100)), cells were incubated with an antibody against RNA polymerase II phosphorylated at serine 5 (RNAPII-S5P; Cell Signaling Technology, cat no. 13523) diluted 1:50 in wash buffer supplemented with 0.1% BSA and 2 mM EDTA for 1 h at room temperature and subsequently overnight at 4 °C. Beads were washed and incubated with anti-rabbit secondary antibody (antibodies-online, cat no. ABIN101961) diluted 1:100 in wash buffer for 1 h at room temperature. After two washes with Triton-300 wash buffer (20 mM HEPES pH 7.5, 300 mM NaCl, 0.5 mM spermidine, 0.1% Triton X-100, protease inhibitor cocktail (Sigma, P8340, 1:100)), the beads were resuspended in Triton-300 wash buffer containing pAG-Tn5 transposase (Cell Signaling Technology) diluted 1:20 and incubated for 1 h at room temperature. Beads were subsequently washed three times with Triton-300 wash buffer and subjected to tagmentation by incubation in 50 µl CUTAC-DMF tagmentation buffer (5 mM MgCl_2_, 10 mM TAPS pH 8.0, 0.05% Triton X-100, and 10% N,N-dimethylformamide) for 20 min at 37 °C. Following tagmentation, the beads were washed with TAPS wash buffer (10 mM TAPS pH 8.5, 0.2 mM EDTA) and resuspended in 5 µl SDS-Proteinase K release solution (0.2% SDS, 1 mg/ml Proteinase K). Samples were incubated at 58 °C for 1 h, followed by addition of 15 µl 0.67% Triton X-100 to neutralize the SDS. Sequencing libraries were amplified using barcoded i5 and i7 primers together with NEBNext 2x PCR Master Mix (New England Biolabs; M0541). Libraries were purified using AMPure XP Reagent (Beckman Coulter) and eluted in 0.1 x TE buffer. Library quality and concentration were assessed using an Agilent TapeStation system. Libraries were sequenced on the NextSeq 2000 platform (Illumina) (paired-end, 2 x 60 bp).

### Immunofluorescence and high-content microscopy

Cells were plated in 96-well plates (Revvity). Immediately before fixation, cells were labeled with 10 µM EdU (Sigma) for 30 min. Cells were fixed with 3% paraformaldehyde and 2% sucrose in PBS for 10 min at room temperature. Cells were permeabilized using 0.2% Triton X-100 (Sigma) in PBS for 5 min and blocked with 3% BSA in PBS for 30 min. Detection of EdU-labeled DNA was performed by copper(I)-catalyzed azide-alkyne cycloaddition in 43 mM Tris pH 7.4, 129 mM NaCl, 4 mM CuSO_4_, 20 µM AFDye 488 Azide (Jena Bioscience), 10 mM L-ascorbic acid for 30 min at room temperature. Next, cells were incubated with the primary antibodies in blocking buffer overnight at 4 °C. After washing, cells were incubated with appropriate fluorophore-conjugated secondary antibodies for 1h at RT. Counterstaining of nuclei was performed by 2.5 mg/ml Hoechst 33342 (Perkin Elmer) for 10 min at RT. Images were acquired with an Operetta CLS High-Content Imaging System (Revvity) at 20x or 40x magnification using water immersion objectives. Images were processed and analyzed using Harmony High-Content Imaging and Analysis Software (Revvity) and custom scripts in R. For the pulse chase, cells were pulse-labeled with EdU for 20 min, washed three times with PBS, and incubated in fresh medium containing 50 ng/ml nocodazole for the indicated chase times before fixation. Fixation and staining were performed as described above. Primary antibodies are listed in Supplementary Table S1.

### MTT assay

Cells were seeded in 96-well plates and treated the following day with either DMSO or 1 µM 5-Ph-IAA and the indicated concentration of prexasertib. 48 h after treatment, 20 µl of thiazolyl blue tetrazolium bromide (5 mg/ml) was added directly to the medium. After 2 h at 37°C, 100 μl DMSO was added, the plate was incubated for 20 minutes at room temperature and the absorbance was measured in a microplate reader at 590 nm. The absorbance was normalized to the negative control of medium without cells, and the cell viability was calculated relative to DMSO control.

### Bioinformatics

For RNA-sequencing, reads were mapped to the Homo sapiens genome hg38 using HISAT2 (2.2.1) ^49^ and counted with featureCounts (2.0.8) ^50^. Lowly expressed genes were filtered prior to analysis, and library sizes were normalized using the trimmed mean of M-values (TMM) method. Differential gene expression analysis was performed using the limma-voom workflow (v3.58.1) with empirical Bayes moderation, and P values were adjusted for multiple testing using the Benjamini-Hochberg method ^51,52^. Fast gene set enrichment analysis (fGSEA, v1.8.0) using C2 and hallmark gene sets from the Molecular Signature Database (MSigDB) was used to identify significantly enriched pathways ^53^. CUTAC reads were mapped to the Homo sapiens genome hg38 using Bowtie2 ^54^. CUTAC peaks were called with MACS2 ^55^. Peaks were called separately for each experimental condition using MACS2. Peak sets from all conditions were concatenated, sorted, and merged using BEDTools ^56^. Read counts within consensus peaks were quantified using featureCounts ^50^. Low-abundance peaks were filtered prior to analysis, and read counts were normalized using the trimmed mean of M-values (TMM) method. Differential chromatin accessibility was assessed using limma-voom with empirical Bayes moderation, and P values were adjusted for multiple testing using the Benjamini-Hochberg method. Peaks were annotated to genomic features and the nearest genes using ChIPseeker ^57^. For promoter accessibility analyses, only peaks located within ±1 kb of annotated transcription start sites were retained.

For integration with RNA-seq data, promoter-associated peaks assigned to the same gene were summarized by calculating the mean log₂ fold change. Genes were classified as showing increased or decreased promoter accessibility when at least one promoter-associated peak showed an adjusted P value < 0.05 and an absolute log₂ fold change ≥ 1 in the corresponding direction. BigWig files were created with deepTools2 ^58^. The Integrative Genomics Viewer (IGV) was used to visualize bigWig files ^59^.

### Statistical Analysis

Statistical methods and the numbers of biological replicates are reported in the corresponding figure legends. Statistical analyses were performed using Prism 10 (GraphPad) and R.

## DATA AVAILABILITY

RNA-sequencing and CUTAC datasets are available at the NCBI’s Gene Expression Omnibus under accession numbers GSE337682 and GSE337683.

## ACKNOWLEDGEMENTS

This work was supported by grants from the Deutsche Forschungsgemeinschaft (DFG, German Research Foundation) GA 575/10-1+2 (to SG) and 440766788 (INST 93/1023-1-FUGG, Operetta CLS system). We thank Marie Zoller and Julia Merk for their help in generating the degron cell line and performing RT-qPCRs.

## AUTHOR CONTRIBUTIONS

DBM, DG, MR, NG, KMA, LD and AB performed the experiments. CSV performed high-content microscopy. CPA performed next generation sequencing. SG performed bioinformatic analysis. ME provided resources. SG planned the study, supervised experiments and wrote the paper.

## DECLARATION OF GENERATIVE AI AND AI-ASSISTED TECHNOLOGIES IN THE WRITING PROCESS

During the preparation of this work the authors used ChatGPT to improve language, grammar, and readability. The authors reviewed and edited the content as needed and take full responsibility for the content of the published article.

