## Supplemental Material for "Replication-coupled chromatin assembly cooperates with ATR/CHK1 signaling to suppress DNA damage and premature MMB-dependent mitotic transcription"

**Figure S1**

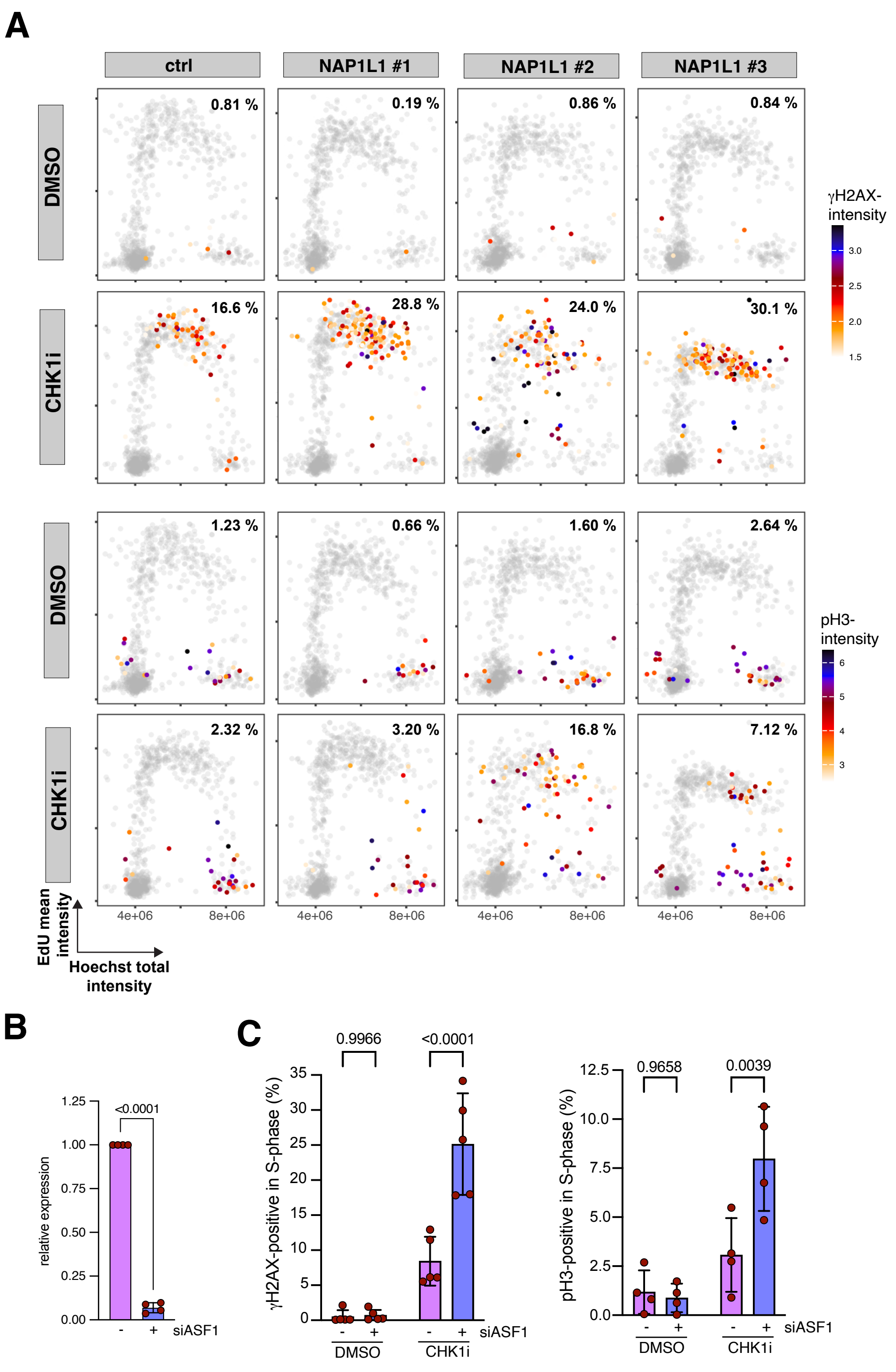


**Figure S1: Depletion of NAP1L1 or ASF1A enhances CHK1i-induced DNA damage and premature mitotic entry.** A) Single-cell analysis of DNA content and DNA synthesis following depletion of NAP1L1 with three independent siRNAs and treatment with CHK1i. DNA content (Hoechst intensity; x-axis) and EdU incorporation (y-axis) are shown for individual cells. Cells are color-coded according to mean γH2AX or phospho-histone H3 (pH3) fluorescence intensity, as indicated. The percentages of γH2AX-positive and pH3-positive S-phase cells are shown. B) Validation of ASF1A depletion by RT-qPCR. Data represent mean ± SD (n = 4). Statistical significance was determined using a two-tailed Student's t-test. C) Quantification of γH2AX-positive and pH3-positive S-phase cells following ASF1 depletion and CHK1 inhibition. Data are shown as mean ± SD. Two-way ANOVA (n = 5).

### Figure S2

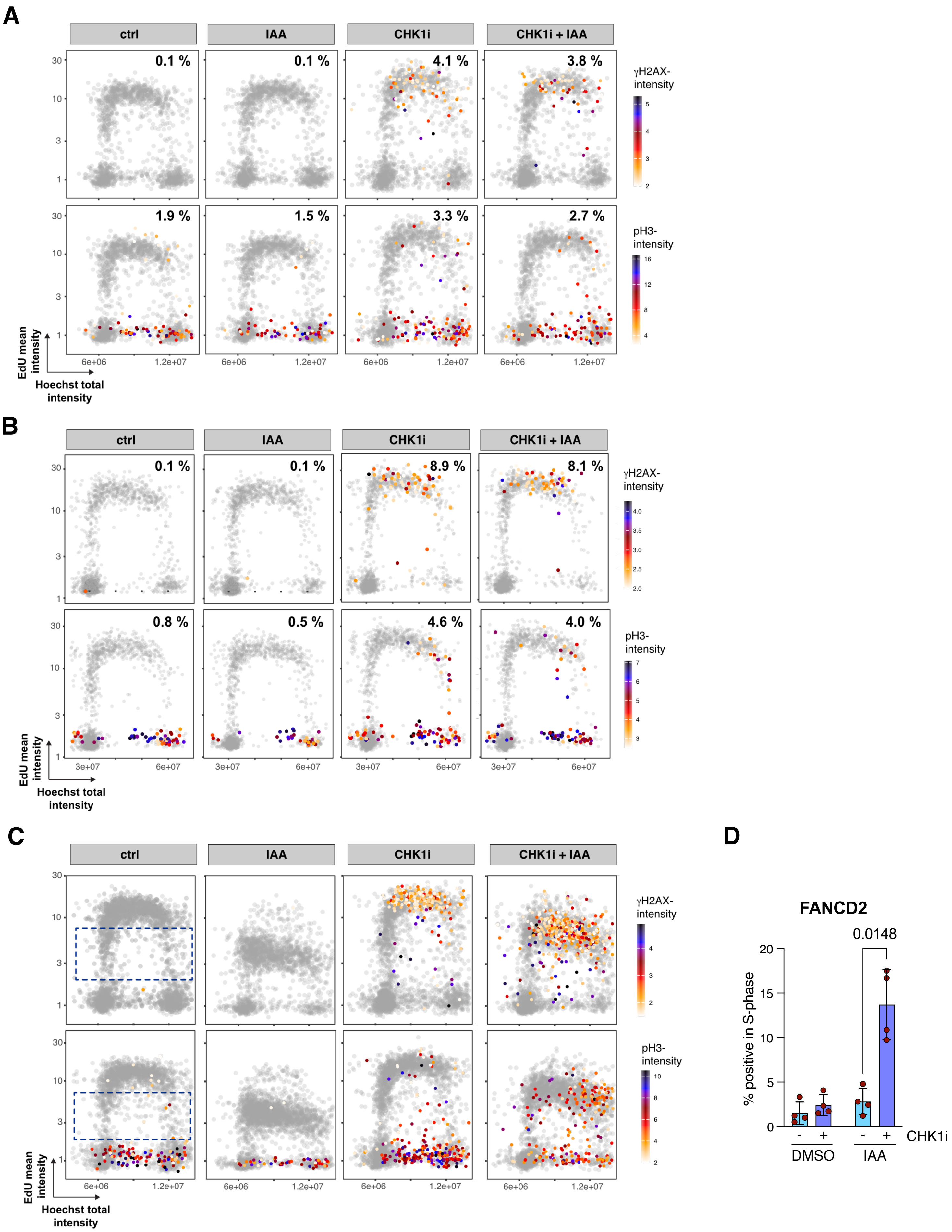
A
ctrl
IAA
CHK1i
CHK1i + IAA
0.1 %
0.1 %
4.1 %
3.8 %
1.9 %
1.5 %
3.3 %
2.7 %
γH2AX-intensity
pH3-intensity
EdU mean intensity
Hoechst total intensity
6e+06
1.2e+07
B
ctrl
IAA
CHK1i
CHK1i + IAA
0.1 %
0.1 %
8.9 %
8.1 %
0.8 %
0.5 %
4.6 %
4.0 %
γH2AX-intensity
pH3-intensity
EdU mean intensity
Hoechst total intensity
3e+07
6e+07
C
ctrl
IAA
CHK1i
CHK1i + IAA
γH2AX-intensity
pH3-intensity
EdU mean intensity
Hoechst total intensity
6e+06
1.2e+07
D
FANCD2
0.0148
% positive in S-phase
CHK1i
DMSO
IAA

**Figure S2: Validation of CHAF1A degron specificity and analysis of DNA damage following CHAF1A depletion and CHK1 inhibition.** A, B) Parental HCT116 (A) and A549 (B) cells expressing TIR1-F74G but lacking the CHAF1A-mAID allele were treated with 5-Ph-IAA and CHK1i as indicated. The percentages of γH2AX-positive and phospho-histone H3 (pH3)-positive S-phase cells were quantified. Data represent two independent experiments. C) HCT116-CHAF1A-mAID cells were treated with 5-Ph-IAA and CHK1i as indicated and analyzed for γH2AX, EdU, and DNA content. Scatter plots show EdU incorporation (y-axis) versus DNA content (x-axis) for individual cells, color-coded according to γH2AX fluorescence intensity. Combined CHAF1A depletion and CHK1 inhibition preferentially increased γH2AX accumulation in a subset of S-phase cells with reduced EdU incorporation. A representative experiment is shown. D) Quantification of S-phase cells positive for FANCD2 following CHAF1A degradation and CHK1 inhibition in A549-CHAF1A-mAID cells (n = 3 independent replicates). Statistical significance was assessed by one-way ANOVA.

### Figure S3

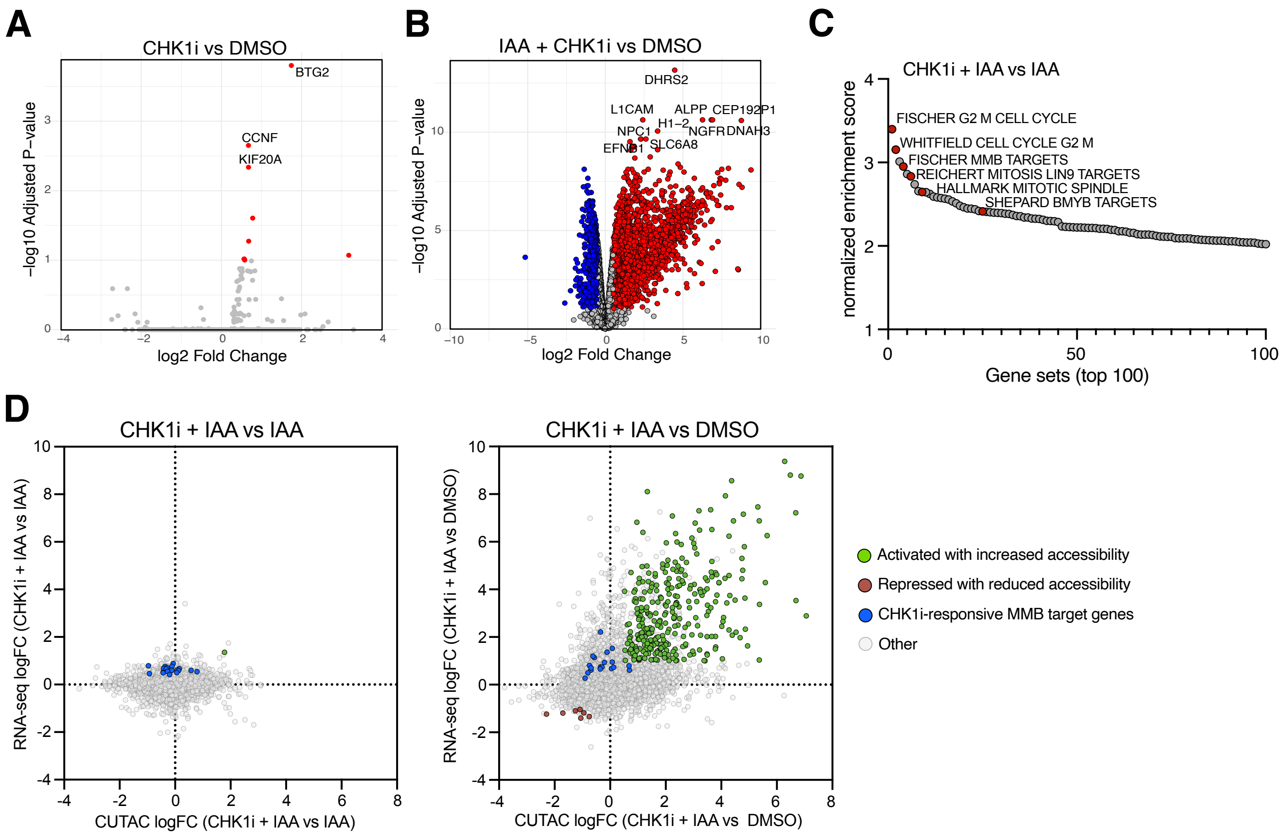


**Figure S3: Transcriptomic and CUTAC-based accessibility analyses following CHAF1A depletion and CHK1 inhibition.** A, B) Volcano plots showing differential gene expression following CHK1 inhibition alone relative to untreated cells (A) or following combined CHAF1A depletion and CHK1 inhibition relative to untreated cells (B). Significantly regulated genes (absolute log2 fold change > 0.5, adjusted $P < 0.1$) are highlighted in red (upregulated) and blue (downregulated). C) Ranked fGSEA results showing the top 100 positively enriched gene sets following CHK1 inhibition in CHAF1A-depleted cells. Gene sets associated with G2/M progression, mitosis, and cell-cycle regulation are highlighted. D) Relationship between changes in gene expression and promoter accessibility following CHK1 inhibition in CHAF1A-depleted cells. Left: Scatter plot showing RNA-seq log2 fold changes and corresponding changes in promoter-associated CUTAC signal within ±1 kb of the transcription start site following CHK1 inhibition in CHAF1A-depleted cells (IAA+CHK1i versus IAA). Right: Scatter plot showing RNA-seq log2 fold changes and corresponding changes in promoter-associated CUTAC signal following combined CHAF1A depletion and CHK1 inhibition relative to untreated cells (IAA+CHK1i versus DMSO). Each point represents one gene. Genes showing concordant increases in gene expression and promoter accessibility are highlighted in green, whereas genes showing concordant decreases in gene expression and promoter accessibility are highlighted in red. CHK1i-responsive MMB target genes defined in Figure 4E are highlighted in blue.

#### Figure S4

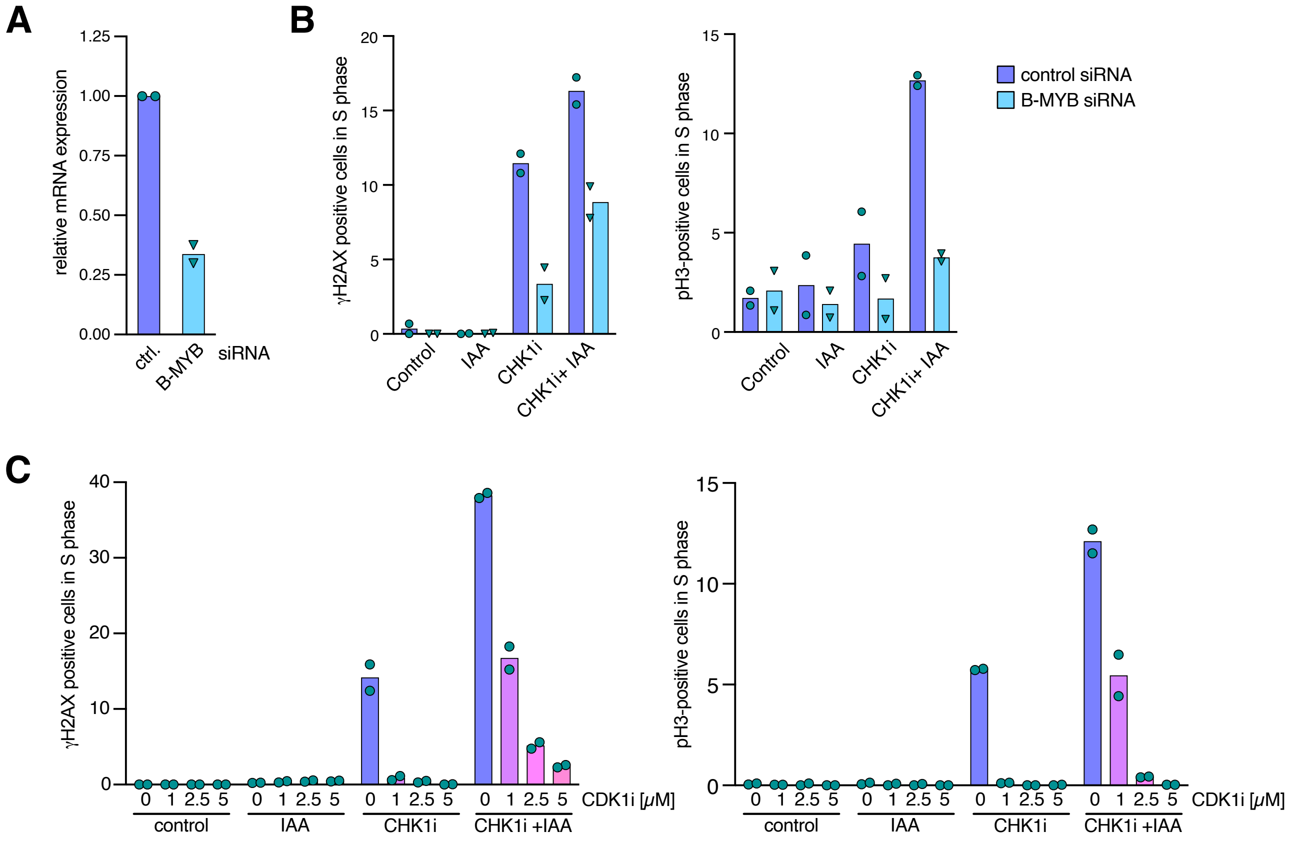


**Figure S4: B-MYB and CDK1 contribute to checkpoint inhibitor sensitivity following CHAF1A loss**
A) Validation of B-MYB depletion by RT-qPCR (n = 2). B) Quantification of γH2AX-positive and phospho-histone H3 (pH3)-positive S-phase cells following siRNA-mediated depletion of B-MYB in HCT116 CHAF1A-mAID cells treated with 5-Ph-IAA and CHK1i as indicated (n = 2 independent experiments). C) Quantification of γH2AX-positive and phospho-histone H3 (pH3)-positive S-phase cells following treatment of A549 CHAF1A-mAID cells with 5-Ph-IAA and CHK1i and with increasing concentrations of the CDK1 inhibitor RO-3306 as indicated (n = 2 independent experiments).

**Supplementary Table S1**

**Antibodies**

| target | company | cat# |
|---|---|---|
| rabbit monoclonal anti-RNAPII-S5P | Cell Signaling Technology | 13523 |
| anti-rabbit secondary antibody | Antibodies Online | ABIN101961 |
| rabbit monoclonal anti-CHAF1A (D77D5) | Cell Signaling Technology | 5480S |
| Mouse monoclonal anti-beta-Actin | Santa Cruz Biotechnology | sc-47778 |
| Rabbit polyclonal anti-pH3 (Ser10) | Millipore | 06-570 |
| Mouse monoclonal anti-pH2A.X (Ser139) | Santa Cruz Biotechnology | sc-517348 |
| Rabbit polyclonal anti-pKAP1 (S824) | Abcam | ab70369 |
| Mouse monoclonal anti-B-MYB (clone LX015.1) | gift from Roger Watson | N/A |
| rabbit polyclonal anti-FANCD2 | Novus Biological | NB100-182 |

**RT-qPCR primers**

| Gene | fw primer | bw primer |
|---|---|---|
| ASF1A | GAGTGCATCGAGGACCTGTC | AACATATGCCTTCCTGCGGG |
| CDCA8 | GCTGACAGCAAAGAGATCTTCC | GCCAATAATCGCAGGCTCT |
| DHRS2 | ACCAGTGAGCAGATCTGGGA | TCCATGTAGGGCAGCAACTG |
| FAM83D | TTGATTGATGGCATCCGCGT | TTCAACCACTTGGCCAGACA |
| GAS2L3 | GCTGTCGGCATGAAGAGC | AATCGATGAGAACAACTACAAGGA |
| KIF14 | CCTGTCTTTTTGCTTATGGTCAG | TCTTCACTAAATCCCATCATCG |
| KIF20A | CGGCGACTAGGTGTGAGTAAG | GGATCCCTTGCGACATGA |
| MYBL2 | CCTGCCCCTATGTCCAGT | CGTGACACCCTCAACACCT |
| PLK1 | AAGATCTGGAGGTGAAAATAGGG | AGGAGTCCCACACAGGGTCT |

**siRNAs**

| Gene | Name/ sequence | company, catalog # |
|---|---|---|
| ctrl | non-targeting control | Thermo Fisher Scientific, 4390843 |
| CHAF1A #1 | GCCUGAAUCUUGUCCCAAAtt | Thermo Fisher Scientific, s19499 |
| CHAF1A #2 | GAAGAAGACUCUGUACUCAtt | Thermo Fisher Scientific, s19500 |
| CHAF1A #3 | CGAAACUUGUCAACGGGAAtt | Thermo Fisher Scientific, s19501 |
| CHAF1B#1 | GGACGGUUACUGCUCAUUUtt | Thermo Fisher Scientific, s15705 |
| CHAF1B#2 | CGAGUAUACAGUAUACAGAtt | Thermo Fisher Scientific, s15704 |
| CHAF1B#3 | GGAUCUGGAAGGUAGAAAAtt | Thermo Fisher Scientific, s15706 |
| NAP1L1#1 | CUUUUUACGUGAGCGUAUAtt | Thermo Fisher Scientific, s9265 |
| NAP1L1#2 | GGAACACGAUGAACCUAUUtt | Thermo Fisher Scientific, s9266 |
| NAP1L1#3 | GCCUCUAUUUGAUAAGCGAtt | Thermo Fisher Scientific, s9267 |
| ASF1A | GCAGAGAGCAGUAAUCCAAtt | Thermo Fisher Scientific, s226043 |
| B-MYB | CCACATCGAAGGAACAGGAtt | Thermo Fisher Scientific, s9117 |
